# The biosynthetic pathway of ubiquinone contributes to pathogenicity of *Francisella*

**DOI:** 10.1101/2021.08.03.455006

**Authors:** Katayoun Kazemzadeh, Mahmoud Hajj Chehade, Gautier Hourdoir, Camille Brunet, Yvan Caspar, Laurent Loiseau, Frederic Barras, Fabien Pierrel, Ludovic Pelosi

**Affiliations:** CNRS, CHU Grenoble Alpes, Grenoble INP, TIMC, Université Grenoble Alpes, Grenoble, France; Laboratoire de Bactériologie-Hygiène Hospitalière, Centre National de Référence des Francisella, Centre Hospitalier Universitaire Grenoble Alpes, 38000, Grenoble, France; Univ. Grenoble Alpes, CHU Grenoble Alpes, CEA, CNRS, IBS, F-38000 Grenoble, France; Aix Marseille Université, CNRS, Laboratoire Chimie Bactérienne, Institut Microbiologie de la Méditerranée, 31 Chemin Joseph Aiguier, Marseille 13009, France; SAMe Unit, Department of Microbiology, Institut Pasteur, Paris, France; IMM-UMR 2001 CNRS-Institut Pasteur, Paris, France

**Keywords:** ubiquinone biosynthesis, coenzyme Q, quinone, aerobic respiration, *Francisella tularensis*, *Francisella novicida*, respiratory chain, metabolism, pathogenicity

## Abstract

*Francisella tularensis* is the causative agent of tularemia. Because of its extreme infectivity and high mortality rate, this pathogen was classified as a biothreat agent. *Francisella* spp are strict aerobe and ubiquinone (UQ) has been previously identified in these bacteria. While the UQ biosynthetic pathways were extensively studied in *Escherichia coli* allowing the identification of fifteen Ubi-proteins to date, little is known about *Francisella* spp. In this study, and using *Francisella novicida* as a surrogate organism, we first identified UQ_8_ as the major quinone found in the membranes of this bacterium. Then, we characterized the UQ biosynthetic pathway in *F. novicida* using a combination of bioinformatics, genetics and biochemical approaches. Our analysis disclosed the presence in *Francisella* of ten putative Ubi-proteins and we confirmed eight of them by heterologous complementation in *E. coli*. The UQ biosynthetic pathways from *F. novicida* and *E. coli* share a similar pattern. However, differences were highlighted: the decarboxylase remains unidentified in *Francisella* spp and homologs of the Ubi-proteins involved in the O_2_-independent UQ pathway are not present. This is in agreement with the strictly aerobic niche of this bacterium. Then, *via* two approaches, i.e. the use of an inhibitor (3-amino-4-hydroxybenzoic acid) and a transposon mutant, which both strongly impair the synthesis of UQ, we demonstrated that UQ is essential for the growth of *F. novicida* in a respiratory medium and contributes to its pathogenicity in *Galleria mellonella* used as an alternative animal model.

**Importance:** *Francisella tularensis* is the causative bacterium of tularemia and is classified as a biothreat agent. Using multidisciplinary approaches, we investigated the ubiquinone (UQ) biosynthetic pathway that operates in *F. novicida* used as a surrogate. We showed that UQ_8_ is the major quinone identified in the membranes of *Francisella novicida*. We identified a new competitive inhibitor, which strongly decreased the biosynthesis of UQ. Our demonstration of the crucial role of UQ for the respiratory metabolism of *F. novicida* and for the involving in its pathogenicity in the *Galleria mellonella* model should stimulate the search for selective inhibitors of bacterial UQ biosynthesis.

## Introduction

*Francisella tularensis* is a Gram-negative, strictly aerobic, facultative intracellular pathogen responsible for tularemia. Infection can occur by inhalation, ingestion, transmission from arthropod vectors or exposure to infected animals (1). After its entry into macrophages, the bacteria are sequestered into phagosomes and prevent further endosomal maturation. *Francisella* cells then disrupt the phagosome and are released into the cytosol in which they rapidly proliferate (2). Eventually, the infected cells undergo apoptosis or pyroptosis, and the progeny bacteria are released to initiate new rounds of infection (2). Currently, there is no suitable vaccine against tularemia and due to its extreme infectivity and high virulence, *F. tularensis* species have been classified as a biothreat agent (3). The genus *Francisella* includes the three species: *F. tularensis, F. novicida* and *F. philomiragia* (4). Moreover, *F. tularensis* is further divided into the subspecies *tularensis* (Type A strains) and *holarctica* (Type B strains), which are the most virulent strains responsible for human disease, whereas *F. philomiragia* and *F. novicida* are avirulent in healthy humans (4). A *F. novicida* type strain U112 is commonly used as a surrogate for *Francisella tularensis* in virulence studies using animal models (5).

The development of genome-scale genetic methods allowed the identification of hundreds of genes participating to variable extents to *Francisella* virulence (6). However, the specific contribution of only a limited number of these genes was demonstrated at the molecular level. Although an important proportion of the identified genes are related to metabolic functions, the relationship between metabolism and the life cycle of *Francisella* is still poorly understood. However, global analysis of genes essential for the growth in culture of *F. novicida* U112 (7) and more recently of that of *F. tularensis* ssp *tularensis* Schu S4 (8) highlighted the involvement of several ubiquitous pathways found in proteobacteria. Among the most significant are the folate pathway, the heme synthesis pathway, the methylerythritol phosphate pathway involved in isoprenoid synthesis, the chorismate pathway and the ubiquinone (UQ) synthesis pathway, on which this work is focused.

Isoprenoid quinones are conserved in most respiratory and photosynthetic organisms and function primarily as electron and proton carriers in the electron-transfer chains. Quinones are composed of a polar redox-active head group linked to a lipid side chain, which varies in both length and degree of saturation (9). Proteobacteria contain two main types of quinone, i.e. benzoquinones and naphthoquinones, represented by UQ (or coenzyme Q) and menaquinone (MK)/ demethylmenaquinone (DMK), respectively (9). UQ is the major electron carrier used for the reduction of dioxygen by various cytochrome oxidases, whereas MK and DMK function predominantly in anaerobic respiratory chains (9). However, as demonstrated recently in *Pseudomonas aeruginosa*, UQ can also be produced and used as a main respiratory quinone under anaerobic conditions (10). Besides its role in bioenergetics, UQ was also reported to be involved in gene regulation, oxidative stress, virulence and resistance to antibiotics (11, 12). More recently, new functions for UQ in bacteria were discovered such as its requirement for *Escherichia coli* to grow on medium containing long chain fatty acids as a carbon source (13). UQ biosynthesis in aerobic conditions has been widely studied in *E. coli* (14). The classical UQ biosynthetic pathway requires twelve proteins (UbiA to UbiK and UbiX). UbiC catalyzes the first committed step in the biosynthesis of UQ, the conversion of chorismate to the 4-hydroxybenzoate (4HB) precursor. Then, UbiA, UbiD to UbiI and UbiX catalyze the prenylation, decarboxylation, hydroxylations and methylations of the phenyl ring of the 4HB to synthesize UQ. In addition, UbiB and UbiK are accessory proteins while UbiJ is involved in the assembly and/or the stability of the aerobic Ubi complex, which was recently characterized in *E. coli* (15). This latter is able to synthesize UQ also under anoxic conditions and we identified three proteins, UbiU, UbiV and UbiT, which are required for UQ biosynthesis only under anoxic conditions (16).

Here, we show that UQ_8_ is the major quinone of *F. novicida* U112. We identified candidate Ubi-proteins in *F. novicida* U112 and validated their functions by heterologous complementation in *E. coli* mutant strains. Our results show that UQ biosynthesis in *Francisella* spp is mostly similar to that of *E. coli*, with the notable absence of UbiX and UbiD for the decarboxylation step. Genetic and chemical inactivation of UQ biosynthesis thanks to a transposon mutant and to a new inhibitor (3-amino-4-hydroxybenzoic acid), respectively, demonstrated that UQ_8_ is crucial for the growth of *F. novicida* in respiratory media and that UQ deficiency impairs the pathogenicity of *F. novicida* against *Galleria mellonella*. Altogether, our results shed light on the role of UQ in the life cycle of *Francisella* and show that UQ contributes to its pathogenicity.

## Results

### UQ_8_ is the major quinone of *F. novicida*

The quinone content of *F. novicida* grown under ambient air at 37°C in Chamberlain media supplemented with either glucose (fermentative medium) or succinate (respiratory medium) as the only carbon source was determined and compared with that of *E. coli* MG1655 grown in the same fermentative medium. In the electrochromatograms of lipid extracts from *F. novicida*, a single peak was observed around 8.3 min, the same retention time as UQ_8_ in *E. coli* extracts (Fig. 1A). Note that in these analyses, UQ_10_ was used as an internal standard, which was added to the samples. MS analysis of the major peak in *F. novicida* extracts showed a predominant ammonium adduct (M^+^ NH_4_^+^) with an m/z ratio of 744.5, together with minor adducts, such as Na^+^ (749.7) and H^+^ (727.8) (Fig. 1B). These masses identify UQ_8_ (monoisotopic mass, 726.5) as the major quinone produced by *F. novicida*. Interestingly, the carbon source in the culture media did not greatly affect the UQ_8_ content (Fig. 1A). The *F. novicida* extracts did not contain any naphthoquinones, unlike *E. coli*, which showed predominantly demethylmenaquinone (DMK_8_) eluting around 12 min. The absence of detectable levels of naphthoquinones in *F. novicida* lipid extracts (Fig. 1A) is in agreement with the absence of menaquinone biosynthesis (Men or futalosine) encoding genes in its genome. Together, our results establish that *E. coli* and *F. novicida* share UQ_8_ as a main quinone in aerobic conditions.

**Figure 1.**
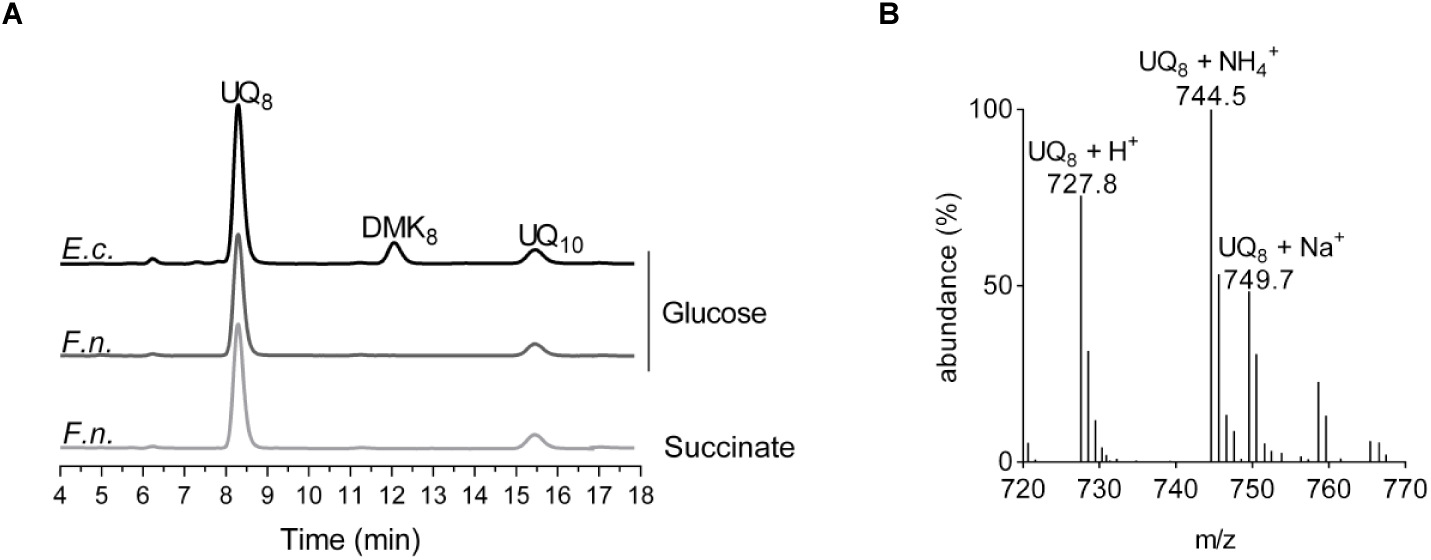
UQ_8_ is the major quinone used by *F. novicida*. **(A)** HPLC-ECD analysis of lipid extracts from 1 mg of *E. coli* MG1655 (*E*.*c*.) and *F. novicida* (*F*.*n*.) cells grown aerobically in Chamberlain medium with 0.4% (w/v) either glucose or succinate as the sole carbon source. The chromatograms are representative of three independent experiments. The peaks corresponding to UQ_8_, DMK_8_ and the UQ_10_ standard are indicated. **(B)** Mass spectrum of the quinone eluting at 8.30 min from extracts of *F. novicida* grown in the Chamberlain media. H^+^, NH_4_^+^ and Na^+^ adducts of UQ_8_ are indicated.

### Identification of Ubi-proteins in the *Francisella* spp

To identify candidate Ubi-proteins in *F. novicida*, UbiX, and UbiA to UbiK from *E. coli* MG1655 were screened for homologs in the protein sequence data set, available at MicroScope (www.genoscope.cns.fr/agc/microscope), using the BLASTP software. As listed in Table S1, this analysis identified eight homologous proteins in *F. novicida*, i.e. UbiA to UbiC, UbiE, UbiG to UbiI and UbiK, called hereafter UbiA_*Fn*_ to UbiC_*Fn*_, UbiE_*Fn*_, UbiG_*Fn*_ to UbiI_*Fn*_ and UbiK_*Fn*_, respectively. Genes *ubiA*_*Fn*_ and *ubiC*_*Fn*_ in one hand and genes *ubiI*_*Fn*_ and *ubiH*_*Fn*_ in another hand present an organization similar to the *ubiC*-*ubiA and the ubiH*-*ubiI* operons from *E. coli*, respectively (12). As reported previously for *Pseudomonas aeruginosa* (17) and *Xanthomonas campestris* (18), *F. novicida* possesses a Coq7 hydroxylase, which is a functional homolog of the UbiF protein found in *E. coli* and other species (19). The detection of an homolog for *E. coli* UbiJ required less restrictive Blast parameters. We noticed that the gene coding for the putative UbiJ candidate *FTN_0460*, hereafter called *ubiJ*_*Fn*_, lies between *ubiE*_*Fn*_ and *ubiB*_*Fn*_, an organization similar to the *ubiE*-*ubiJ*-*ubiB* operon from *E. coli* (20). UbiJ_*Fn*_ has 21% amino-acid identity with UbiJ from *E. coli* and both proteins contain a sterol carrier protein 2 domain in their N-terminal regions(http://pfam.xfam.org/) (21). The same Ubi-proteins were identified in the highly virulent strain *F. tularensis* ssp *tularensis* Schu S4 (Table S1).

Homologs of UbiD and UbiX were not yet identified and the counterparts of these two proteins in *Francisella* spp remain to be determined. The work is in progress in our laboratory. Under anaerobic conditions, *E. coli* still synthesizes UQ, and we recently identified three genes, which we called *ubiT, ubiU*, and *ubiV*, as essential for this process (16). Homologs of *ubiT, ubiU*, and *ubiV*, which participate to the O_2_-independent UQ biosynthetic pathway, were not identified in the screened *Francisella* genomes (Table S1), in agreement with the strictly aerobic metabolism of *Francisella* spp. In overall, these data show that the O_2_-dependent UQ biosynthetic pathways in *F. novicida*, in *F. tularensis* and in *E. coli* are related, the major difference being the absence of UbiX-UbiD for the decarboxylation step (Fig. S1).

### Functional characterization of *Ubi*_*Fn*_*-*proteins in *E. coli*

To test whether the candidate Ubi-proteins identified in *F. novicida* were indeed involved in UQ biosynthesis, we assessed their capacity to functionally complement *E. coli* strains in which the UQ protein-encoding genes were inactivated (Δ*ubiAc*, Δ*ubiBc*, Δ*ubiCc*, Δ*ubiEc*, Δ*ubiFc*, Δ*ubiGc*, Δ*ubiHc*, Δ*ubiIc*, Δ*ubiJ* and Δ*ubiKc*, see Table S2). We assessed the quinone content and the capacity to grow on solid minimal medium containing fermentable (glucose) or respiratory (succinate) carbon sources. *E. coli* Δ*ubiAc*, Δ*ubiBc*, Δ*ubiGc*, Δ*ubiHc* and Δ*ubiJ* transformed with empty vector are unable to synthesize UQ_8_ (Fig. 2A) and are thus unable to grow on a respiratory medium (Fig. 2C). In contrast, their growth on a fermentative medium is not affected (Fig. 2C). Except for the Δ*ubiAc* mutant strain, in which the prenylation reaction of the 4HB is impaired, most mutants accumulate an early intermediate corresponding to octaprenylphenol (OPP) (Fig. 2A and S1). *E. coli* Δ*ubiEc* and Δ*ubiFc* cells accumulate C2-demethyl-C6-demethoxy-UQ_8_ (DDMQ_8_) and C6-demethoxy-UQ_8_ (DMQ_8_), which are the substrates of UbiE and UbiF, respectively (Fig. 2A and S1). We found that UbiA_*Fn*_, UbiB_*Fn*_, UbiE_*Fn*_, Coq7_*Fn*_ and UbiG_*Fn*_ restored growth of *E. coli* Δ*ubiAc*, Δ*ubiBc*, Δ*ubiEc*, Δ*ubiFc* and Δ*ubiGc* cells on respiratory medium (Fig. 2C) and allowed for UQ_8_ biosynthesis in LB medium to 96, 26, 7, 49 and 38% of the level of UQ_8_ present in the WT cells, respectively (Fig. 2A and 3A). Concomitantly, OPP content decreased and Coq7_*Fn*_ abolished the accumulation of DMQ_8_ in Δ*ubiFc* cells (Fig. 2A). As we previously reported, *E. coli* Δ*ubiIc* and Δ*ubiKc* cells displayed a strong decrease in UQ_8_ (22, 23), but the residual UQ_8_ content was sufficient to support growth on succinate (Fig. 2A and 2C). Similar results were obtained with Δ*ubiCc* cells grown in minimal M9 medium (Fig. 2B and 2C), which had to be used instead of LB since the later contains 4HB that restores normal UQ_8_ content in Δ*ubiCc* (data not shown). In all three strains, the expression of the corresponding Ubi proteins, UbiC_*Fn*_, UbiI_*Fn*_ *and* UbiK_*Fn*_, increased significantly the UQ_8_ content (Fig. 3A and 3B). Since the increase obtained in Δ*ubiIc* cells was moderate (from 25 to 40%), we further confirmed the ability of UbiI_*Fn*_ to catalyze *C5-hydroxylation* by using an *E. coli* Δ*ubiIc*Δ*ubiF* strain. This deletion mutant lacks C5- and C6-hydroxylation activities and consequently accumulates 3-octaprenyl-4-hydroxyphenol (4HP_8_) (22). We found that UbiI_*Fn*_ was able to restore DMQ_8_ biosynthesis in *E. coli* Δ*ubiIc*Δ*ubiF* cells (Fig. S2), i.e. to catalyze C5 hydroxylation, concomitantly to a strong decrease of 4HP_8_. Taken together, all these results confirm unambiguously that UbiA_*Fn*_, UbiB_*Fn*_, UbiC_*Fn*_, Coq7_*Fn*_, UbiE_*Fn*_, UbiG_*Fn*_, UbiI_*Fn*_ and UbiK_*Fn*_ are the functional counterpart of the *E. coli* Ubi-proteins and we propose that they compose the biosynthetic pathway of UQ_8_ *in F. novicida*. Only two proteins, UbiJ_*Fn*_ and UbiH_*Fn*_ did not complement the *E. coli* Δ*ubiJ* and Δ*ubiHc* (Fig. 2A, 2C and 3A). The low percentage of identity between UbiJ and UbiH from *E. coli* and their homologs in *F. novicida* (21 and 27%, respectively) could explain these results (Table S1).

**Figure 2.**
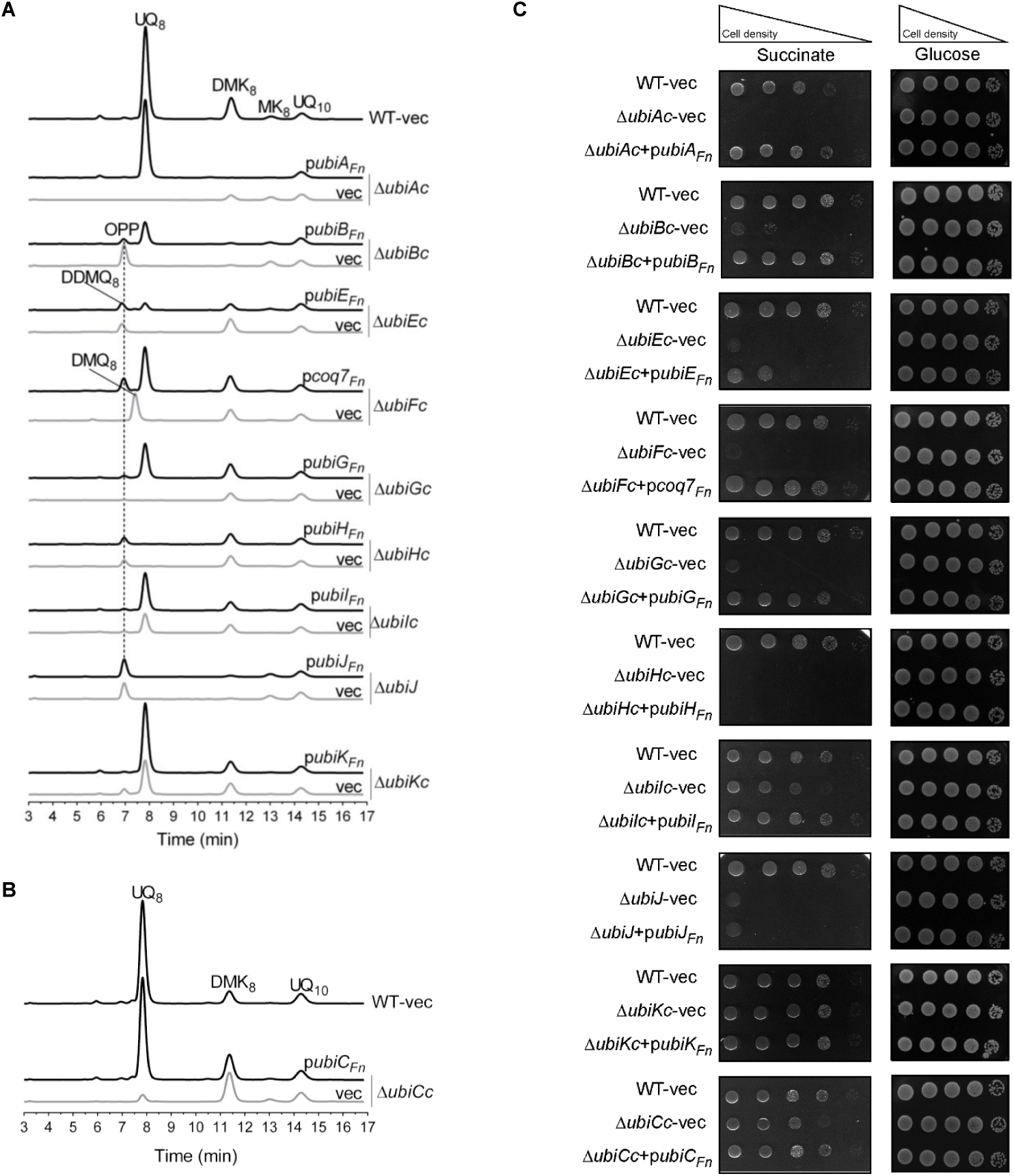
Complementation analysis of *E. coli* UQ_8_ biosynthesis mutants with the putative Ubi-proteins from *F. novicida*. The Δ*ubi E. coli* mutant strains transformed with pTrc99a (vec) or pTrc99a encompassing the *ubi*_*Fn*_ genes were grown over night at 37°C in LB medium **(A)** or in M9 minimal medium **(B)** with 0.4% (w/v) glucose as the sole carbon source. Expression of the Ubi_*Fn*_-proteins was induced by addition of IPTG to a final concentration of 100 µM. The *E. coli* wild-type strain MG1655 (WT) transformed with the pTrc99a empty vector was used as a control. HPLC-ECD analysis of lipid extracts from 1 mg of cells. The chromatograms are representative of three independent experiments. The peaks corresponding to OPP, DDMQ_8,_ DMQ_8_, UQ_8_, MK_8_, DMK_8_ and the UQ_10_ standard are indicated. **(C)** Serial dilutions were spotted onto plates containing M9 minimal medium with 0.4% (w/v) either glucose or succinate as the sole carbon source and IPTG (100 µM final concentration). The plates were incubated overnight at 37°C.

**Figure 3.**
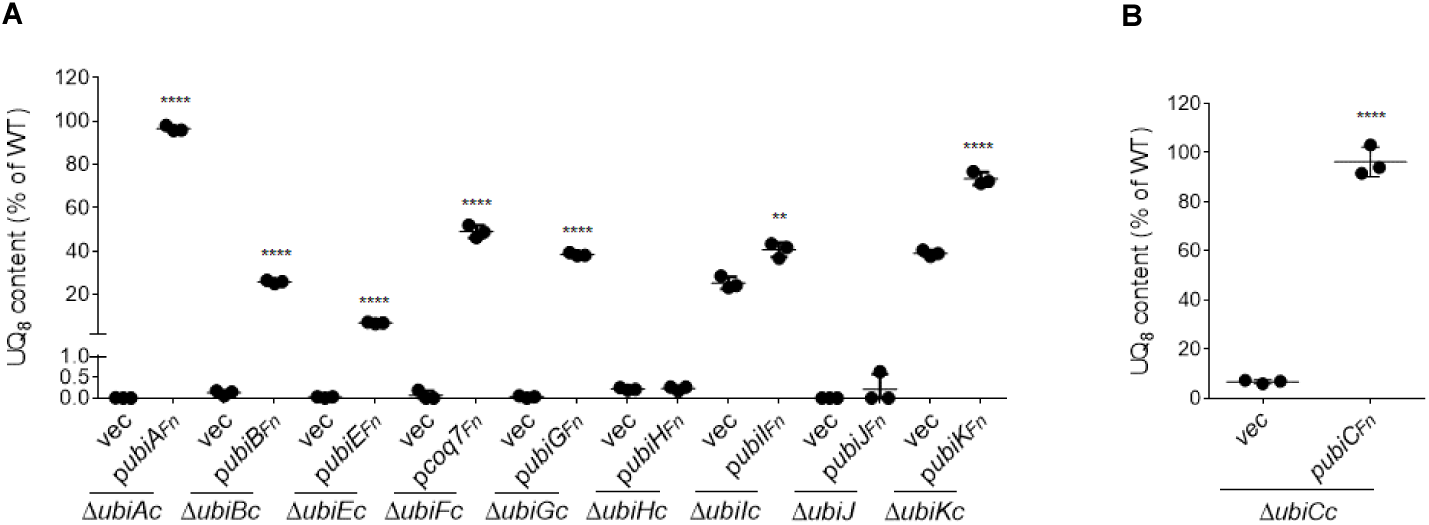
Quantification of cellular UQ_8_ contents of Δ*ubi E. coli* mutant strains expressing the Ubi_*Fn*_-proteins. The Δ*ubi E. coli* mutant strains transformed with pTrc99a (vec) or pTrc99a encompassing the *ubi*_*Fn*_-genes were grown over night at 37°C in LB medium **(A)** or in M9 minimal medium **(B)** with 0.4% (w/v) glucose as the sole carbon source. Expression of the Ubi_*Fn*_-proteins was described in the legend of the Figure 2. Quantifications are expressed as the percentages of the control value, which corresponds to the UQ_8_ content of the wild-type strain (n = 3). ****, P < 0.0001 and **, P < 0.005 by unpaired Student’s t test.

### UQ_8_ biosynthesis is essential for the growth of *F. novicida* in respiratory medium

To evaluate the physiological importance of UQ for *F. novicida*, we screened *ubi* genes in the *F. novicida* transposon (Tn) mutant library available at the Manoil Laboratory (7). Only the Tn mutant of *ubiC*_*Fn*_ (called hereafter Tn*-ubiC*_*Fn*_) was available in the library, and we compared this mutant strain to its isogenic control strain U112 (Table S2). Recall that UbiC catalyzes the first committed step in the biosynthesis of UQ, i.e. the conversion of chorismate to 4HB (Fig. S1). First, we showed that the growth of Tn*-ubiC*_*Fn*_ cells under ambient air in respiratory Chamberlain medium was severely impaired compared to the WT (Fig. 4A). In contrast, the growth of *F. novicida* in fermentative medium was less affected (Fig. 4B). In parallel, UQ_8_ content was strongly lowered in Tn*-ubiC*_*Fn*_ cells from 166 to 7 pmol/mg cells in fermentative medium and from 134 to 11 pmol/mg cells in respiratory medium (Fig. 4C). As expected, addition of 4HB to the culture rescued the growth of Tn *ubiC*_*Fn*_ in respiratory medium and increased the UQ_8_ content to WT levels (Fig. 4C). Taken together, these results show the overall requirement of UQ_8_ for the growth of *F. novicida* especially in respiratory medium.

**Figure 4.**
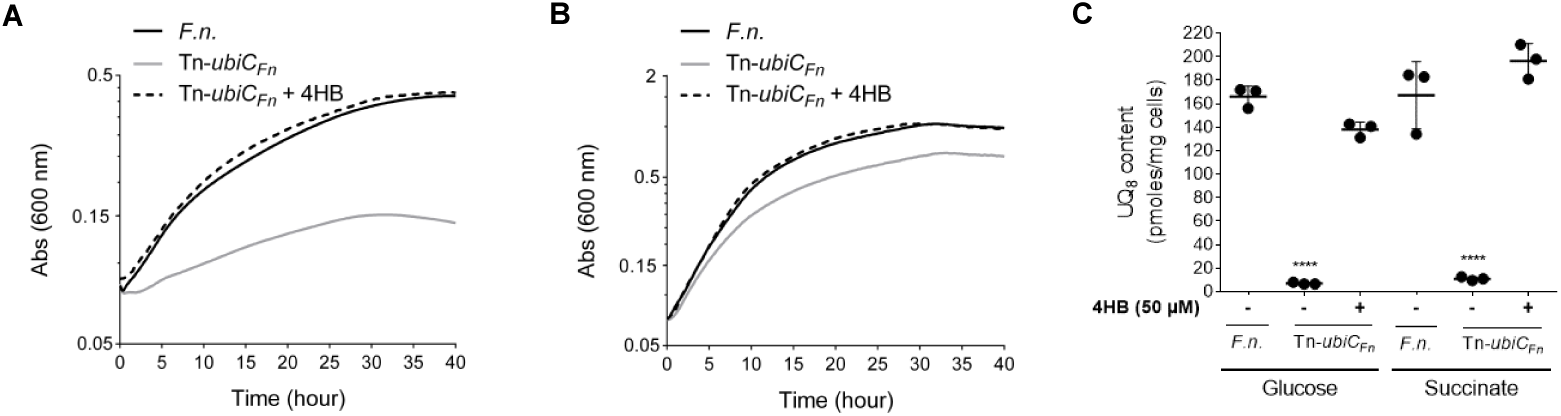
UQ_8_ is essential for the growth of *F. novicida* in respiratory medium. *F. novicida* (*F*.*n*.) and transposon mutant of *ubiC*_*F*n_ (Tn-*ubiC*_*Fn*_) were grown aerobically in Chamberlain medium with 0.4% (w/v) either succinate **(A)** or glucose **(B)** as the sole carbon source. Growth (average of sixtiplicate growth curves) was followed as the change in the absorbance at 600 nm in a Tecan plate reader. **(C)** Cellular UQ_8_ contents were quantified for *F*.*n*. and Tn-*ubiC*_*Fn*_ according to the materials and methods section. 4HB was added to rescue the growth and the UQ_8_ biosynthesis in Tn-*ubiC*_*Fn*_. Quantifications are expressed as pmoles per mg of cells (n = 3). ****, P < 0.0001 by unpaired Student’s t test.

### 3A4HB inhibits UQ_8_ biosynthesis and impairs the growth of *F. novicida* in respiratory medium

Besides genetic inactivation of the UQ pathway, we were interested in the possibility to decrease UQ levels by chemical inhibition. Since we had found UQ to be particularly important for growth of *F. novicida* in respiratory medium (Fig. 4A), we screened for compounds that could inhibit growth in such medium. We tested several compounds: 3 amino-4-hydroxybenzoic acid (3A4HB), 4-amino-benzoic acid (pABA), 4-amino-2-methoxy benzoic acid (pA2MBA) and 4-amino-3-methoxy-benzoic acid (pA3MBA). All these molecules are analogs of 4HB, the native precursor of UQ (Fig. S3A). We observed that bacterial growth was slightly affected in respiratory medium in presence of pABA and pA2MBA, while pA3MBA inhibited growth both in fermentative and respiratory media (Fig. S3B-C). Interestingly, 3A4HB strongly impaired bacterial growth in respiratory medium, while inhibition was milder in fermentative medium (Fig. S3B-C). Based on these results, we followed up on this compound.

We then examined how and to what extent 3A4HB could affect UQ_8_ biosynthesis in *F. novicida*. Bacteria were cultured under ambient air in fermentative Chamberlain medium supplemented with 3A4HB (from 10 µM to 1 mM, final concentration). Endogenous UQ_8_ content was measured in the bacterial cells and compared to a control condition in which only DMSO was added. Figure 5A shows that the UQ_8_ content decreased with increasing concentrations of 3A4HB in the medium, with 0.5 mM yielding to a ∼90% decrease of the UQ_8_ content. Concomitantly, we confirmed that growth of *F. novicida* in presence of 1 mM 3A4HB was strongly impaired in respiratory medium (Fig. 5B), but less so in a fermentative medium (Fig. 5C). Control experiments showed that addition of 4HB to the growth medium counteracted the negative effect of 3A4HB, both in terms of UQ_8_ biosynthesis and bacterial growth (Fig. 5A to 5C).

**Figure 5.**
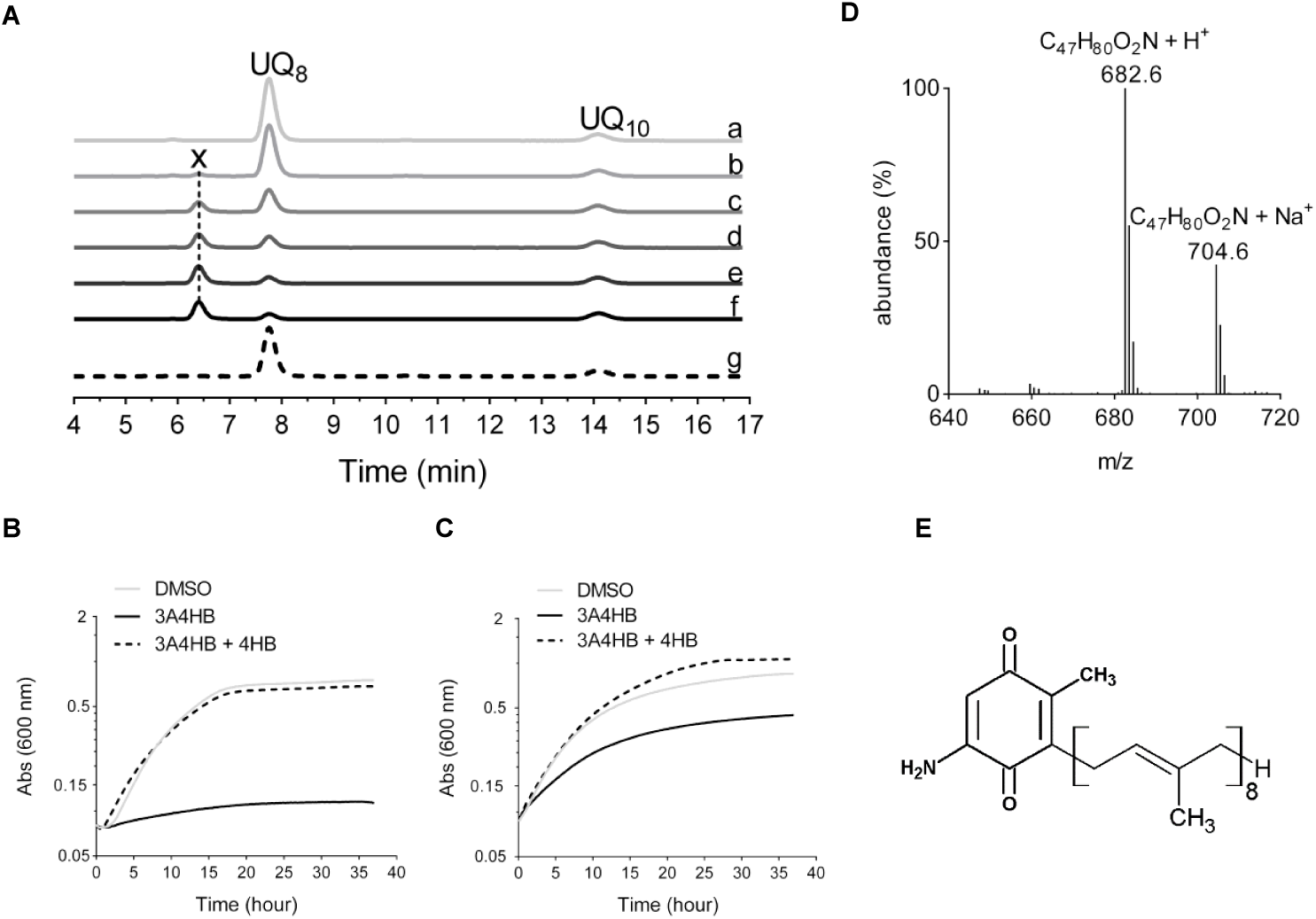
Effect of 3A4HB on UQ_8_ biosynthesis and growth of *F. novicida*. **(A)** HPLC ECD analysis of lipid extracts from 1 mg of *F. novicida* cells grown aerobically in Chamberlain medium with 0.4% (w/v) glucose as the sole carbon source and in the presence of different concentration of 3A4HB solubilized in DMSO (a = DMSO; b = 0.01 mM; c = 0.1 mM; d = 0.25 mM; e = 0.5 mM; f = 1 mM; g = 1 mM 3A4HB + 1 mM 4HB). The chromatograms are representative of three independent experiments. The peaks corresponding to UQ_8_ and the UQ_10_ standard, are indicated. Compound X eluting at 6.5 min is marked. Growth curves for *F. novicida* cultured under aerobic conditions in Chamberlain medium with 0.4% (w/v) either succinate **(B)** or glucose **(C)** as the sole carbon source and in the presence of either DMSO (control), 1 mM 3A4HB or 1 mM 3A4HB + 100 µM 4HB. The growth for each condition (average of sixtiplicate growth curves) was followed as the change in the absorbance at 600 nm in a Tecan plate reader. **(D)** Mass spectrum of compound X eluting from extracts of *F. novicida* grown in the Chamberlain media with 1 mM of 3A4HB. H^+^ and Na^+^ adducts corresponding to this molecule were indicated. **(E)** Proposed structure of compound X in its oxidized form.

Treatment with 3A4HB caused the accumulation of a redox compound that eluted at 6.5 min (compound X on Fig. 5A). MS analysis of this peak showed a predominant proton adduct (M^+^ H^+^) with an m/z ratio of 682.6, together with a minor sodium adduct (M^+^ Na^+^) with an m/z ratio of 704.6 (Fig. 5D). Both species are compatible with a monoisotopic mass of 681.7 g.mol^-1^, which could correspond to that of 2-octaprenyl-3-methyl-6-amino-1,4 benzoquinone (Fig. 5E). According to the sequence of reactions proposed in Figure S1, the formation of compound X would result from prenylation of 3A4HB, decarboxylation and hydroxylation at C1 and then methylation at C3. Thus, 3-octaprenyl-2-methyl-5-amino-1,4 benzoquinone seems to be the “dead-end” product of the UQ_8_ pathway in *F. novicida* cells treated with 3A4HB. Collectively, these results demonstrate unequivocally that 3A4HB acts as a competitive inhibitor of UQ_8_ biosynthesis and affects particularly the respiratory metabolism of *F. novicida*.

### UQ_8_ is involved in the pathogenesis of *F. novicida* in the later steps of the infection

We evaluated the importance of UQ in the pathogenicity of *F. novicida* by studying the Tn*-ubiC*_*Fn*_ mutant. To assess the overall virulence of the Tn*-ubiC*_*Fn*_ strain in a whole organism, we used the wax moth (*G. mellonella*) infection model, which was previously used in studies of human pathogenic and closely related opportunistic and non-pathogenic *Francisella* spp, such as *F. novicida* (24-27). We monitored the survival of larvae infected with the Tn*-ubiC*_*Fn*_ strain or with the isogenic strain U112 as control. When the larvae are turning grey/black and no movement of the larval legs can be observed, they are considered dead (Fig. 6A). The Tn*-ubiC*_*Fn*_ strain was found to be statistically much less virulent than the wild type, but was nevertheless still capable to kill *Galleria* larvae (Fig. 6B). This result suggests that UQ_8_ is involved in the virulence potential of *F. novicida* in *G. mellonella*. To better understand the role of UQ_8_ in the different stages of infection in *G. mellonella*, the pathogenicity of the isogenic control strain pre-treated with 1 mM 3A4HB was studied in order to mimic a UQ_8_ acute deficiency. Recall that this treatment causes a ∼90% decrease of the UQ_8_ content (Fig. 5A) but the inhibition should be alleviated over the infection cycle in the larvae where 3A4HB is not present. The pretreatment with 3A4HB has no effect on the capacity of *F. novicida* to kill *Galleria* larvae (Fig. 6C), suggesting that UQ_8_ does not contribute to the virulence of *Francisella novicida* in the early steps of the infection but more likely in later ones. This result contrasts with that obtained with the Tn*-ubiC*_*Fn*_ strain, which represents a chronic deficiency of UQ_8_.

**Figure 6:**
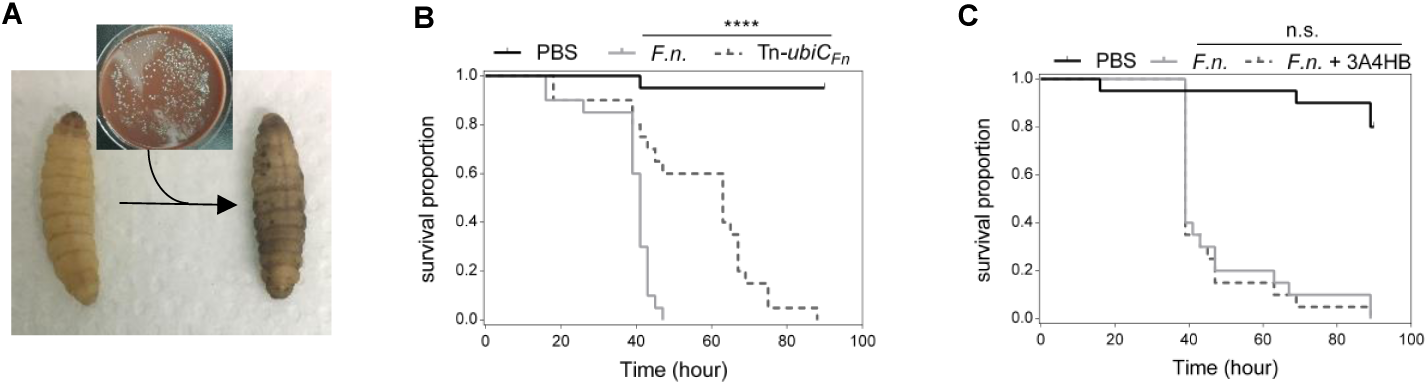
UQ_8_ contributes to the later steps of the infection of *F. novicida* in *G. mellonella*. **(A)** The larvae are turning grey/black when infected. **(B)** Survival curve of *G. mellonella* infected with either *F. novicida* (*F*.*n*.) or the transposon mutant of *ubiC*_*F*n_ (Tn *ubiC*_*Fn*_). ****, P < 0.0001 by Log-rank (Mantel-Cox) test. **(C)** Survival curve of *G. mellonella* infected with *F. novicida* (*F*.*n*.) pre-treated or not with 3A4HB (1 mM final concentration). Each group of *G. mellonella* (n=20) was injected with around 10^6^ CFU/larva and PBS injection was used as control. n.s. (no significant), P = 0.6670 by Log-rank (Mantel-Cox) test.

## Discussion

The chemical analysis performed in this paper established that UQ_8_ is the major isoprenoid quinone synthesized by *F. novicida*. In two representative *Francisella* genomes, we identified homologs for nine of the twelve genes, which are currently known to contribute to UQ biosynthesis in *E. coli* under aerobic conditions. We confirmed the function of seven of the nine homologs by heterologous complementation of *E. coli* Δ*ubi* mutants. From these results, we show again that *E. coli* is a good model to study the function of most exogenous *ubi*-genes (19). We could not confirm the function of UbiH_*Fn*_ and UbiJ_*Fn*_, but the fact that *ubiH*_*Fn*_ and *ubiJ*_*Fn*_ show the same genetic organization as in *E. coli* (a *ubiI*-*ubiH* operon and a *ubiE*-*ubiJ*-*ubiB* operon) strongly supports the implication of these genes in the UQ biosynthetic pathway. Interestingly, both proteins are part of the Ubi-complex in *E. coli* (15). We hypothesize that the low identity of UbiH_*Fn*_ and UbiJ_*Fn*_ with their *E. coli* homologs (∼25%) might impair their assembly within the *E. coli* Ubi-complex and thus compromise our *in vivo* complementation assays. Another possibility relates to the proposed implication in UQ biosynthesis of a non-coding RNA partially overlapping the ORF of UbiJ from *E. coli* (28). We note that the expression of UbiJ from *X. campestris* was also unable to complement an *E. coli* Δ*ubiJ* strain (18). *Francisella* spp shares with *P. aeruginosa* and *X. campestris* a yeast Coq7 protein homolog, which catalyzes the C6-hydroxylation as UbiF from *E. coli* (17, 18, 29). As we demonstrated previously, the Coq7 proteins are found in all three subclasses, alpha-, beta- and gamma-proteobacteria. In contrast, homologs of UbiF proteins are limited to the gamma-proteobacteria (19). Our analysis disclosed also the presence in *Francisella* spp of UbiI and UbiH homologous proteins, which catalyze C5- and C1-hydroxylation in *E. coli*, respectively (22, 30). Consequently, we propose that both *E. coli* and *Francisella* spp share a UQ biosynthetic pathway involving three hydroxylases, i.e. UbiI, UbiH and UbiF in *E. coli* or UbiI, UbiH and Coq7 in *Francisella* spp. Several studies highlighted that the enzymes involved in multiple steps of the UQ biosynthetic pathway vary between bacterial species (14), like for the hydroxylation steps (19), or for the production of 4HB from chorismate by UbiC or XanB2 proteins (31). The decarboxylation step involves UbiD and UbiX in *E. coli* (32), but we could not identify homologs in *Francisella* genomes. A candidate gene *ubiZ* was proposed based on its co-localization with *ubiE* and *ubiB* in the genomes of *Acinetobacter* spp and *Psychrobacter* sp. PRwf-1, which are also devoid of homologs of UbiD and UbiX (33). However, *ubiZ* was not confirmed functionally and this gene is not conserved in *Francisella* genomes. This suggests the existence of another decarboxylation system operating in UQ biosynthesis in *Francisella* spp and potentially in other bacteria lacking *ubiX* and *ubiD* (34).

To assess the essentiality of the UQ biosynthetic pathway in the respiratory metabolism of *F. novicida*, two different approaches were carried out. First, we showed that a transposon mutation of *ubiC*_*Fn*_ gene, which decreases 4HB synthesis, impaired the growth of *F. novicida* mainly in respiratory medium. Interestingly, among all the *ubi*-gene identified in *Francisella* genomes, only *ubiC* was mutated in large-scale studies (7, 8). This supports that the other ubi genes are essential for the viability of *Francisella* spp and strengthens the idea that UQ is key for the development of these bacteria. We noted that the mutation of the *ubiC* gene affects more severely *F. novicida* than *E. coli* for growth in respiratory media, despite both mutants producing comparable amounts of UQ. (∼7-8% compared to the WT) (Fig. 3B and 5A). As *E. coli* synthesizes naphthoquinones but *F. novicida* does not, we propose that the milder phenotype of the *E. coli ubiC* mutant results from naphthoquinones participating to aerobic respiration, as previously suggested (35). Second, we tested the effect of structural analogs of 4HB and we showed that 3A4HB impaired the growth of *F. novicida* mainly in respiratory medium in agreement with a strong decrease of UQ_8_ biosynthesis. We demonstrated that 3A4HB competes with endogenous 4HB and progresses through several steps of the UQ biosynthetic pathway to form the redox compound X that we propose to be 3 octaprenyl-2-methyl-5-amino-1,4-benzoquinone. As the Tn*-ubiC*_*Fn*_ strain and the control strain U112 treated with 1 mM 3A4HB yielded both to a ∼90% decrease of the UQ_8_ content and presented a strong impairment of the growth in a respiratory medium (Fig. 4 and 5), we propose that compound X would not be used as a quinone in the respiratory chain of *F. novicida*.

We noted that homologs of *ubiT, ubiU*, and *ubiV*, which belongs to the O_2_ independent UQ biosynthetic pathway characterized in *E. coli* and *P. aeruginosa* (10, 16), were not identified in the screened genomes of *Francisella* spp. This result is in agreement with the strictly aerobic metabolism of these bacteria. Indeed, tricarboxylic acid (TCA) cycle and the UQ-dependent electron-transfer chain, leading to efficient oxidative phosphorylation, take place in *Francisella* spp (36). A possible link between stress defense and the TCA cycle was previously suggested in *Francisella* pathogenesis (37). Unfortunately, the contribution of UQ and the electron-transfer chain to virulence has not been well documented to date in *Francisella* spp. Using *G. mellonella* as infection model at the scale of an entire organism, we demonstrated through the study of the Tn*-ubiC*_*Fn*_ mutant and the isogenic control strain pre treated with 3A4HB, that UQ_8_ contributes to the virulence *of F. novicida* and more likely in the later steps of the infection, during which the bacteria undergo extensive replication (38). Such a notion support the view that, as other facultative intracellular bacteria, *Francisella* spp are able to use several substrates in order to grow in various environments, such as macrophages. Glycerol via the gluconeogenesis and amino acids were identified as main sources of carbon during intracellular replication of *Francisella* spp in the host cells (36, 39). However, glycerol requires UQ to be efficiently metabolized *via* the ubiquitous enzyme GlpD (40) and amino acids degradation is closely linked to the TCA cycle, which produces reducing equivalents in *Francisella* spp (36). Besides its requirement for bioenergetics, UQ might also contribute to the antioxidant capacity of *Francisella* since it was shown to be a potent lipid soluble antioxidant in *E. coli* (41). During its intracellular life, *Francisella* is exposed to oxidative stress. Indeed, as a defense mechanism for the clearance of phagocytosed microorganisms, both macrophages and neutrophils produce reactive oxygen species, which in turn trigger bacterial killing by causing damage to macromolecules (42, 43). We propose that the reduced content in UQ in the Tn*-ubiC*_*Fn*_ mutant could therefore affect *F. novicida*’s oxidative defense. This hypothesis is in good agreement with recent data showing that reduced expression of UbiC_*Fn*_ decreases the resistance of *F. novicida* to oxidative stress (44). In a similar way, we showed previously that UbiE, UbiJ and UbiB proteins were needed in *Salmonella enterica* serovar *Typhimurium* intracellular proliferation in macrophages (21). Collectively, all these data assign a role for Ubi-proteins in bacterial intracellular proliferation and, more generally, highlight the importance of UQ production for bacterial virulence.

## Materials and methods

### Bacterial strains and growth conditions

All bacterial strains used in this study are listed in Table S2. *F. novicida* U112 was obtained from the Centre National de Référence des *Francisella*, CHU Grenoble-Alpes, France. The transposon mutant Tn*-ubiC*_*Fn*_ in the *F. novicida* U112 strain was obtained from the Manoil Laboratory, Department of Genome Science, University of Washington (7). Both strains were grown on Polyvitex enriched chocolate agar plates (PVX-CHA, bioMérieux, Marcy L’Etoile, France) incubated at 37°C for 48-72 h. Liquid cultures were carried out at 37°C with rotary shaking at 200 rpm in Chamberlain medium (45) supplemented with either glucose or succinate (0.4% (wt/vol) final concentration) as the only carbon source. For growth studies, overnight cultures were used to inoculate a 96-well plate to obtain a starting optical density at 600 nm (OD600) of around 0.1 and further incubated under shaking at 37°C. Changes in OD_600_ were monitored every 10 min for 40 h using the Infinite 200 PRO microplate reader (Tecan, Lyon, France). When required, the medium was supplemented with 4HB in DMSO at 50-100 µM final concentration, pABA, pA2MBA and pA3MBA in DMSO at 1 mM final concentration or 3A4HB at 10 µM-1 mM, final concentration. For CFU counting, bacteria were suspended in PBS and cell suspensions were serially diluted in PBS. For each sample, 100 µL of at least four different dilutions were plated on PVX-CHA plates and incubated for 72 h at 37°C, and CFU were counted using a Scan 100 Interscience.

The *E. coli* Δ*ubiA* and Δ*ubiJ* mutants were constructed as described previously (46). Briefly, the *ubiA::cat* and *ubiJ::cat* mutation was generated in a one-step inactivation of the *ubiA* and *ubiJ* genes. A DNA fragment containing the *cat* gene flanked with the 5’ and 3’ regions of the *ubiA* and *ubiJ* genes was PCR amplified using pKD3 as a template and oligonucleotides 5’-wannerubiA/3’-wannerubiA and 5’-wannerubiJ/3’-wannerubiJ, respectively (Table S3). The Δ*ubiB* mutant was generated as follows: the *cat* gene was inserted in *ubiB* gene between the two sites NruI at 842 and 1004 pb. Then, *ubiB::cat* was PCR amplified using oligonucleotides 5’-xbaIubiB/3’-xbaIubiB (Table S3). Strain BW25113, carrying the pKD46 plasmid, was transformed by electroporation with the amplified fragments and Cat^r^ colonies were selected. The replacement of chromosomal *ubi* by the *cat* gene was verified by PCR amplification in the Cat^r^ clones. *E. coli* K-12 strains JW5713 and JW2226 from the Keio Collection (47) were used as donors in transduction experiments to construct the Δ*ubiC::kan* and Δ*ubiG::kan* mutants of *E. coli* MG1655 strains. The Δ*ubiA*, Δ*ubiB*, Δ*ubiC*, Δ*ubiE*, Δ*ubiG* and Δ*ubiK* strains were cured with pCP20 to yield Δ*ubiAc*, Δ*ubiBc*, Δ*ubiCc*, Δ*ubiEc*, Δ*ubiGc* and Δ*ubiKc* strains, respectively (Table S2). *E. coli* strains (K12, MG1655 or Top10) were grown on lysogeny broth (LB)-rich medium or in M9 minimal medium (supplemented with glucose or succinate, 0.4% (wt/vol) final concentration) at 37°C. Ampicillin (100 µg/ml), kanamycin (50 µg/ml), chloramphenicol (35 µg/ml) and IPTG (100 µM) were added when needed.

### Cloning, plasmid construction, and complementation assays

The plasmids and the primers used in this study are listed in Tables S2 and S3 (supplemental material), respectively. All the plasmids produced in this work were verified by DNA sequencing (GATC Biotech, Konstanz, Germany). The *FTN_0385* (*ubiA*_*Fn*_), *FTN_0459* (*ubiB*_*Fn*_), *FTN_0386* (*ubiC*_*Fn*_), *FTN_0461* (*ubiE*_*Fn*_), *FTN_1146* (*coq7*_*Fn*_), *FTN_0321* (*ubiG*_*Fn*_), *FTN_1237* (*ubiH*_*Fn*_), *FTN_1236* (*ubiI*_*Fn*_), *FTN_0460* (*ubiJ*_*Fn*_) and *FTN_1666* (*ubiK*_*Fn*_) inserts were obtained by PCR amplification using the *F. novicida* U112 genome as template and the oligonucleotides described in Table S3. Inserts were EcoRI-BamHI or EcoRI-HindIII digested and inserted into EcoRI-BamHI- or EcoRI-HindIII-digested pTrc99a plasmids, respectively, yielding the p*ubiA*_*Fn*_, p*ubiB*_*Fn*_, p*ubiC*_*Fn*_, p*ubiE*_*Fn*_, p*coq7*_*Fn*_, p*ubiG*_*Fn*_, p*ubiH*_*Fn*_, p*ubiI*_*Fn*_, p*ubiJ*_*Fn*_ and p*ubiK*_*Fn*_ plasmids (Table S3). The plasmids were transformed into *E. coli* MG1655 strains with mutation of the *ubiA, ubiB, ubiC, ubiE, ubiF, ubiG, ubiH, ubiI, ubiJ* and *ubiK* genes (single and double mutations, see Table S2), and complementation of the UQ_8_ biosynthetic defect was assessed by both measuring the quinone content and plating serial dilutions onto solid M9 minimal medium supplemented with glucose or succinate (0.4% (wt/vol) final concentration) as the only carbon sources and overnight growth at 37°C. Expression of the Ubi-proteins was induced by addition of IPTG to a final concentration of 100 µM.

### Lipid extractions and quinone analysis

Cultures (5 ml under ambient air) were cooled down on ice 30 min before centrifugation at 3200 X g at 4 °C for 10 min. Cell pellets were washed in 1 ml ice-cold PBS and transferred to preweighted 1.5 mL Eppendorf tubes. After centrifugation at 12,000 g at 4 °C for 1 min, the supernatant was discarded, the cell-wet weight was determined (∼5–30 mg) and pellets were stored at -20°C. Quinone extraction from cell pellets was performed as previously described (22). Lipid extracts corresponding to 1 mg of cell-wet weight were analyzed by HPLC electrochemical detection-MS (ECDMS) with a BetaBasic-18 column at a flow rate of 1 mL/min with mobile phases composed of 50% methanol, 40% ethanol, and a mix of 90% isopropanol, 10% ammonium acetate (1 M), and 0.1% TFA). When necessary, MS detection was performed on an MSQ spectrometer (Thermo Scientific) with electrospray ionization in positive mode (probe temperature, 400°C; cone voltage, 80 V). Single-ion monitoring detected the following compounds: UQ_8_ (M^+^ NH4^+^), m/z 744-745, 6–10 min, scan time of 0.2 s; UQ_10_ (M^+^ NH4^+^), m/z 880–881, 10-17 min, scan time of 0.2 s; DMQ_8_ (M^+^ NH4^+^), m/z 714–715–10 min, scan time of 0.4 s; DDMQ_8_ (M^+^ NH4^+^), m/z 700-701, 5-8 min, scan time of 0.4 s; OPP (M^+^ NH4^+^), 656.0-657, 5-9 min, scan time of 0.4 s; compound X (M^+^ H^+^), m/z 682-683, 5-10 min, scan time of 0.4 s. MS spectra were recorded between m/z 600 and 900 with a scan time of 0.3 s. ECD and MS peak areas were corrected for sample loss during extraction on the basis of the recovery of the UQ_10_ internal standard and then were normalized to cell wet weight. The peaks of UQ_8_ obtained with electrochemical detection or MS detection were quantified with a standard curve of UQ_10_ as previously described (22).

### Infections of *G. mellonella* larvae

Larvae of the wax moth *G. mellonella* were purchased from Lombri’carraz SARL, Mery, France. Healthy and uniformly white larvae measuring around 3 cm were selected for infection. The bacteria were grown over night to an OD_600_ of around three. Culture medium was removed by centrifugation and bacteria were diluted in PBS to 10^8^ CFU/mL. Insulin cartridges were sterilized before filling with bacterial solutions. Larvae were injected with 10 μL of bacterial suspensions (10^6^ CFU per larva as recommended (24)) using an insulin pen or with 10 µL of PBS only. The precise number of bacteria transferred in injections was determined by spotting serial dilutions onto chocolate agar plates, and counting CFU after growth at 37°C for 48h. Infected larvae were placed in Petri dishes and maintained at 37°C. Survival of larvae was monitored for 6 days by counting the number of dead larvae each day. A cohort of 20 larvae was used per condition, and the experiment was performed twice. As a control, an untreated cohort of larvae was also followed.

## Acknowledgments

This work was supported by the Agence Nationale de la Recherche (ANR), project O2-taboo ANR-19-CE44-0014, project Emergence (TIMC-UGA), the University Grenoble Alpes (UGA) and the French Centre National de la Recherche Scientifique (CNRS). We thank Dr Patricia Renesto for constructive discussions, technical assistance and Professor Laurent Aussel for critically reading the paper.

## Contributions

F.B., F.P., and L.P. conceived the project and its design. K.K., M.H.C. and G.H. conducted experiments and performed data analysis. K.K., C.B. and Y.C. performed experiments on *G. mellonella*. L.L. contributed to new reagents (strains). All authors edited the manuscript. L.P. wrote the manuscript. L.P. supervised the project.

## Abbreviations

The abbreviations used are:

MK_8_: menaquinone 8;
DMK_8_: dimethyl-menaquinone 8;
OHB: 3 octaprenyl-4-hydroxybenzoic acid;
OPP: octaprenylphenol;
DMQ_8_: C6-demethoxy ubiquinone 8;
DDMQ_8_: C1-demethyl-C6-demethoxy-ubiquinone 8;
UQ_8_: ubiquinone 8;
3A4HB: 3-amino-4-hydroxybenzoic acid;
pABA: 4-amino-benzoic acid, pA2MBA, 4-amino 2-methoxy-benzoic acid and pA3MBA, 4-amino-3-methoxy-benzoic acid;
ECD: electrochemical detection;
Tn: transposon;
OD_600_: optical density at 600 nm and IPTG, isopropyl-1-thio-β-D-galactopyranoside.

---

**Figure S1:**
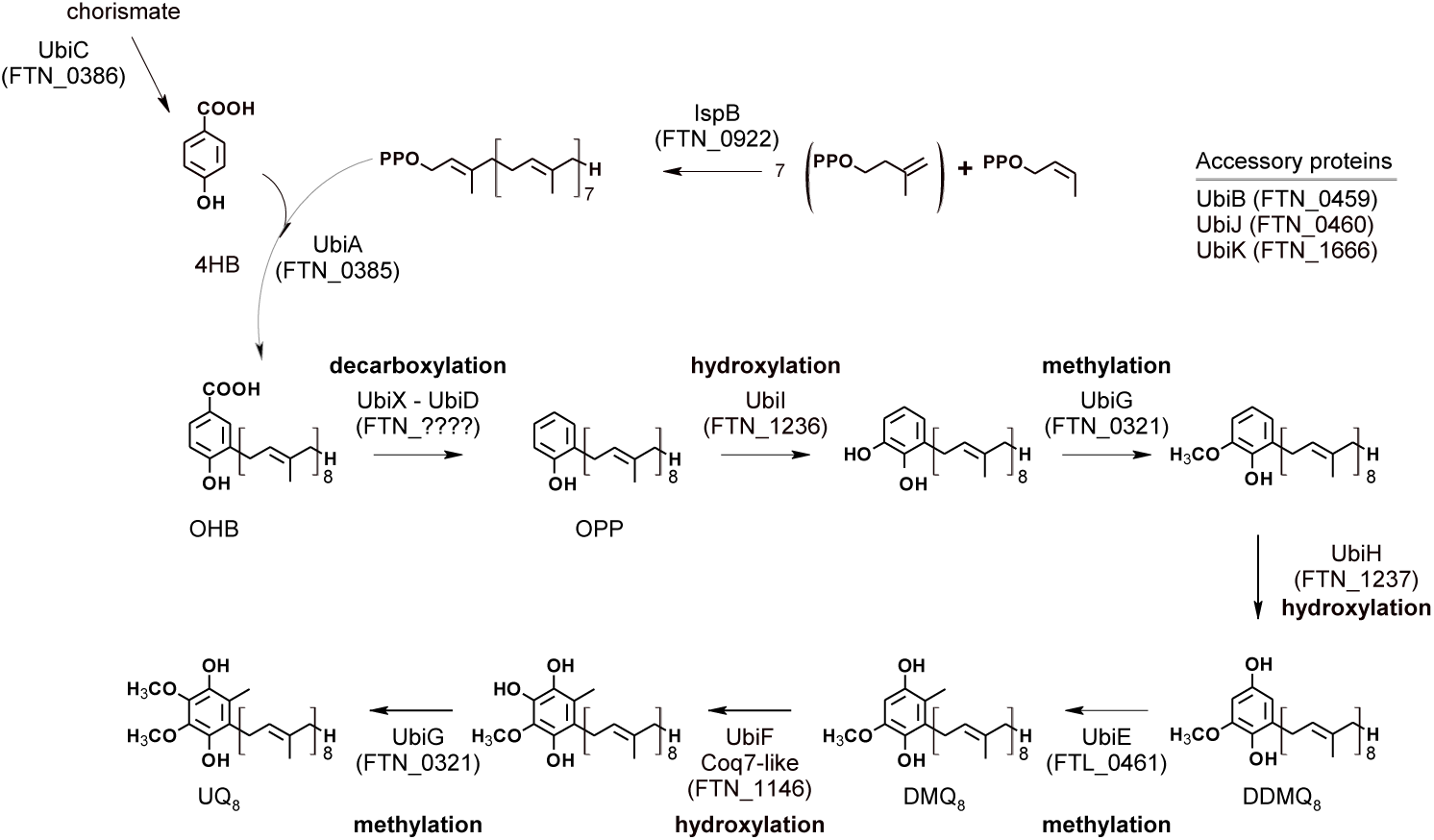
Proposed UQ_8_ biosynthetic pathway in *F. novicida* deduced from the one characterized in *E. coli*. Corresponding protein IDs in *F. novicida* are indicated in parenthesis. There is no identified counterpart of UbiD and UbiX in *F. novicida* proteome. UbiF is only identified in *E. coli* and its functional homolog in *F. novicida* is a Coq7 hydroxylase. Abbreviations used are 4HB, 4-hydroxybenzoic acid; OHB, 3-octaprenyl-4 hydroxybenzoic acid; OPP, octaprenylphenol; DMQ_8_, C6-demethoxy-ubiquinone 8; DDMQ_8_, C1-demethyl-C6-demethoxy-ubiquinone 8; UQ_8_, ubiquinone 8.

**Figure S2.**
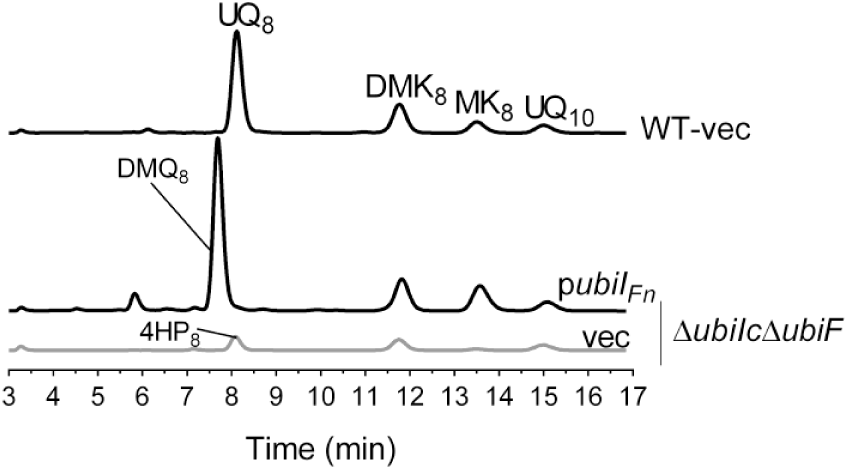
Complementation analysis of *E. coli* Δ*ubiIc*Δ*ubiF* mutant with UbiI_*Fn*>_. The *E. coli* Δ*ubiIc*Δ*ubiF* mutant strain transformed with pTrc99a (vec) or pTrc99a encompassing the *ubiI*_*Fn*_ gene were grown over night at 37°C in LB medium. Expression of the UbiI_*Fn*_ protein was induced by addition of IPTG to a final concentration of 100 µM. The *E. coli* wild-type strain MG1655 (WT) transformed with the pTrc99a empty vector was used as a control. HPLC-ECD analysis of lipid extracts from 1 mg amounts of cells. The chromatograms are representative of three independent experiments. The peaks corresponding to 4HP_8_, 3 octaprenyl-4-hydroxyphenol; DMQ_8_, C6-demethoxy-ubiquinone; UQ_8_, ubiquinone 8; MK_8_, menaquinone 8; DMK_8_, demethylmenaquinone 8 and UQ_10_, ubiquinone 10 as a standard, are indicated.

**Figure S3.**
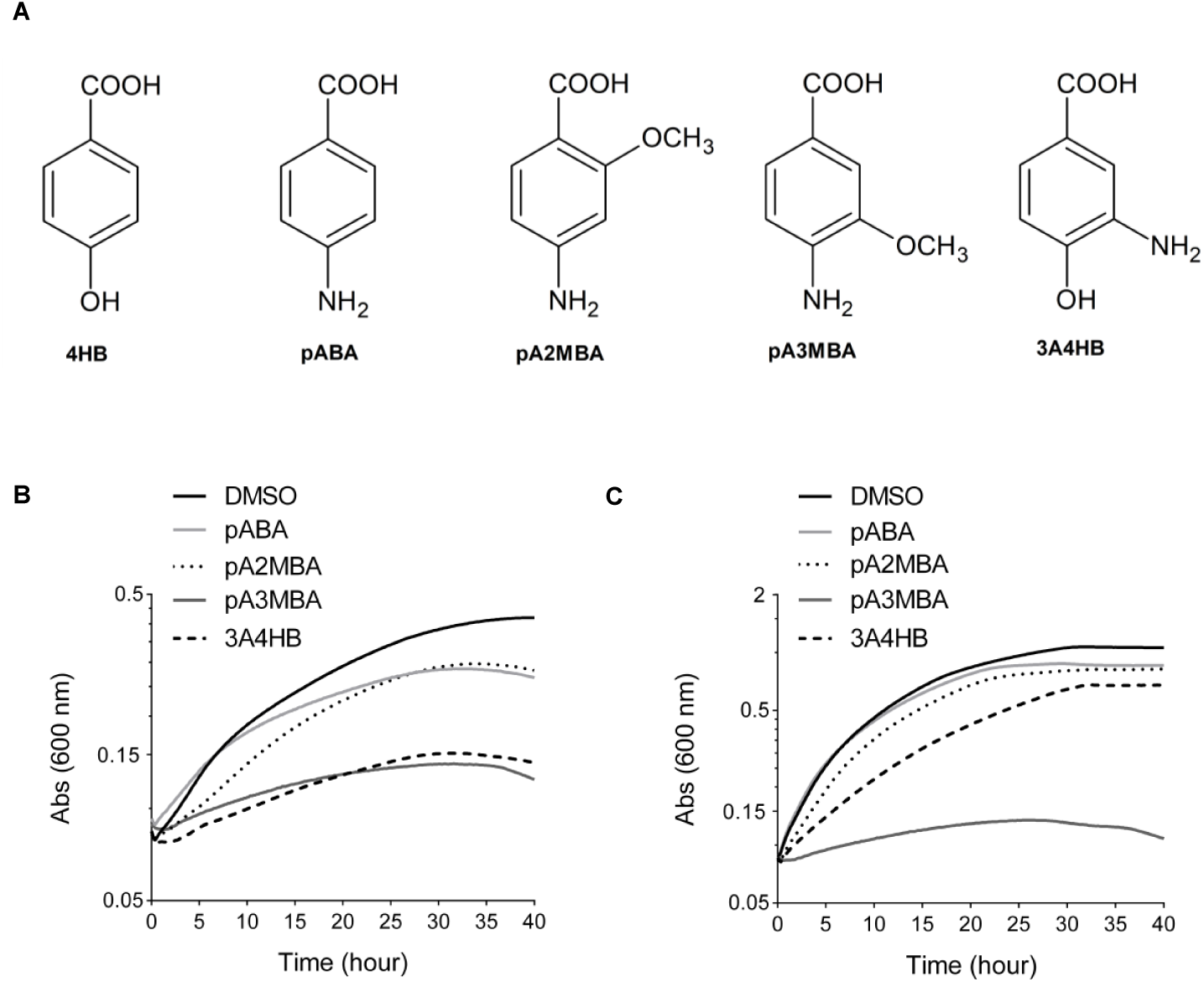
Effect of derivative compounds of 4HB on *F. novicida* growth. **(A)** Chemical structures of the compounds tested. Growth curves for *F. novicida* cultivated under aerobic conditions in Chamberlain medium with 0.4% (w/v) succinate **(B)** or glucose **(C)** as the sole carbon source and in the presence of 1 mM of the various 4HB analogues solubilized in DMSO (average of sixtiplicate growth curves). DMSO was used as a control. The growth for each condition was followed as the change in the absorbance at 600 nm in a Tecan plate reader. More details of growth experiments are found in Materials and methods section. pABA, 4-amino-benzoic acid; pA2MBA, 4-amino-2-methoxy-benzoic acid; pA3MBA, 4 amino-3-methoxy-benzoic acid and 3A4HB, 3-amino-4-hydroxybenzoic acid.

**Table S1:**
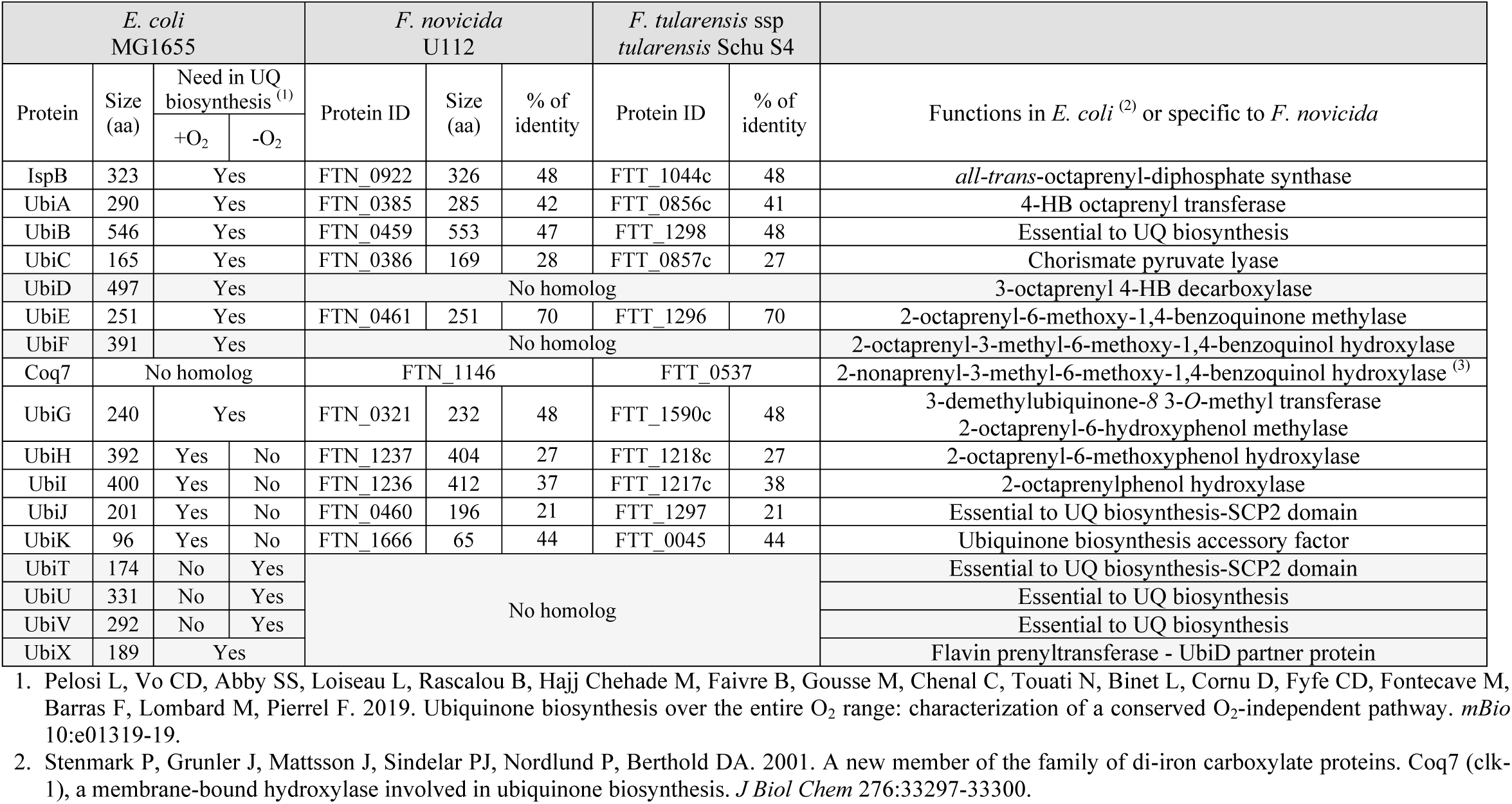
Bioinformatics analysis of the proteins required for UQ biosynthesis in *E. coli* MG1655, in *F. novicida* U112 and in *F. tularensis* ssp *tularensis* Schu S4. ^(1)^ Deduced from (1). ^(2)^ From https://ecocyc.org/. ^(3)^ From (2).

**Table S2:**
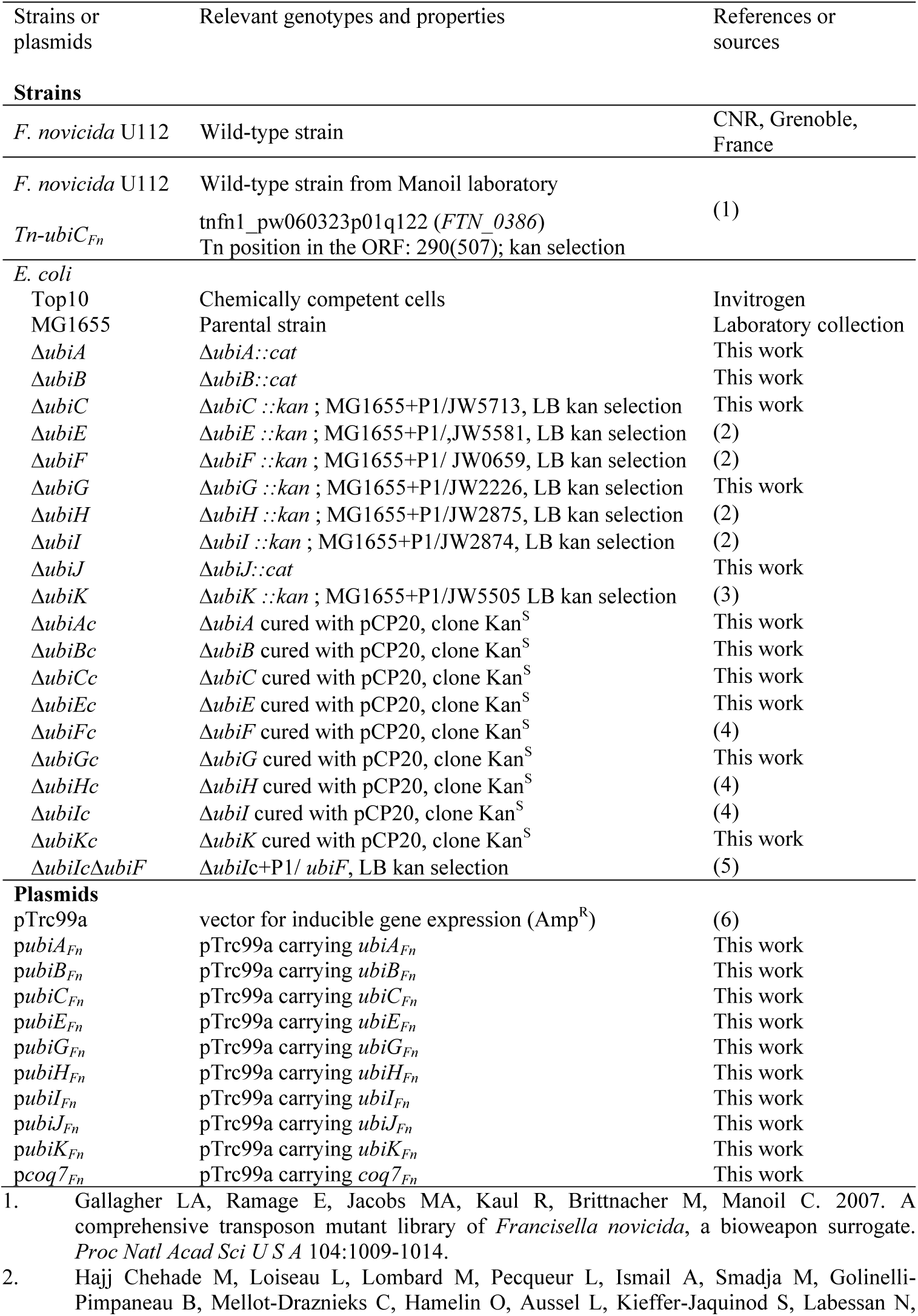
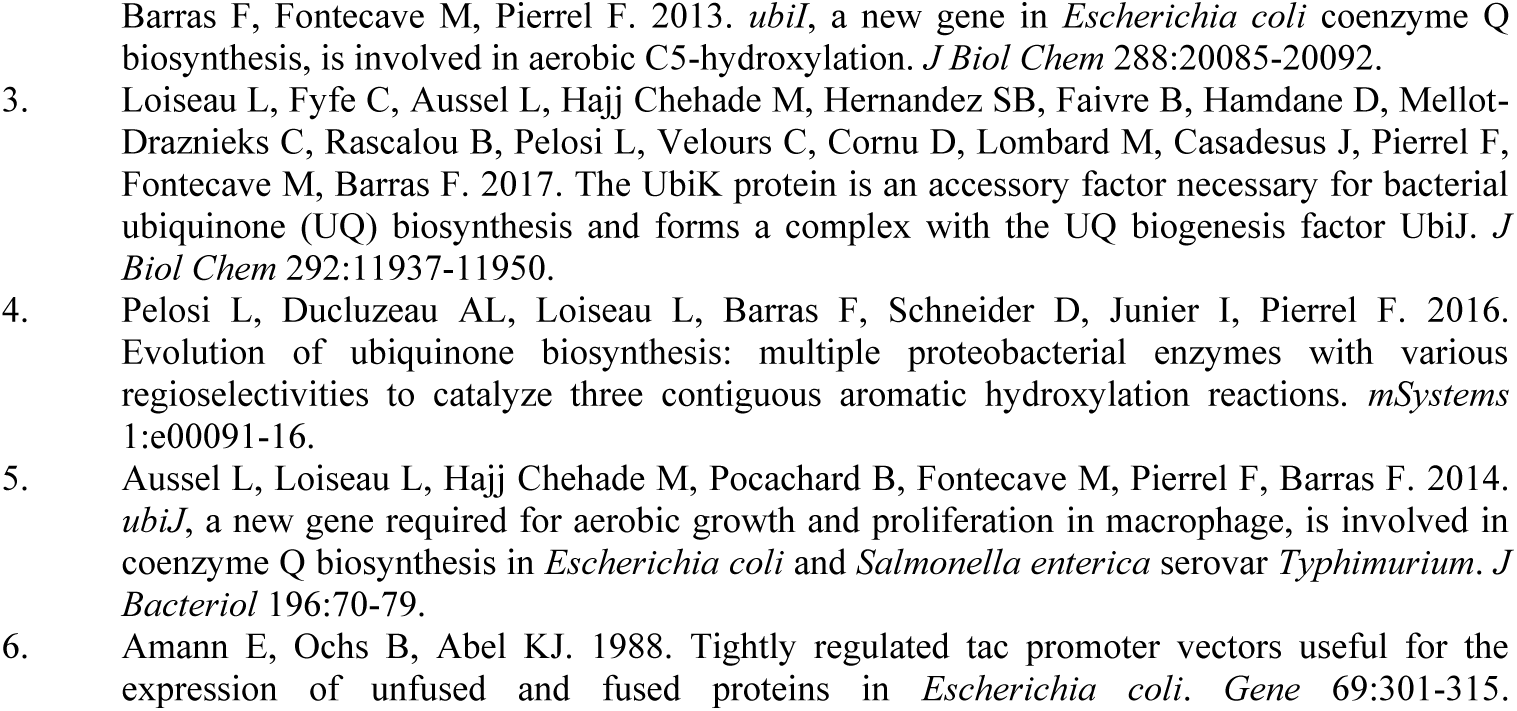
Bacterial strains and plasmids used in this work.

**Table S3:**
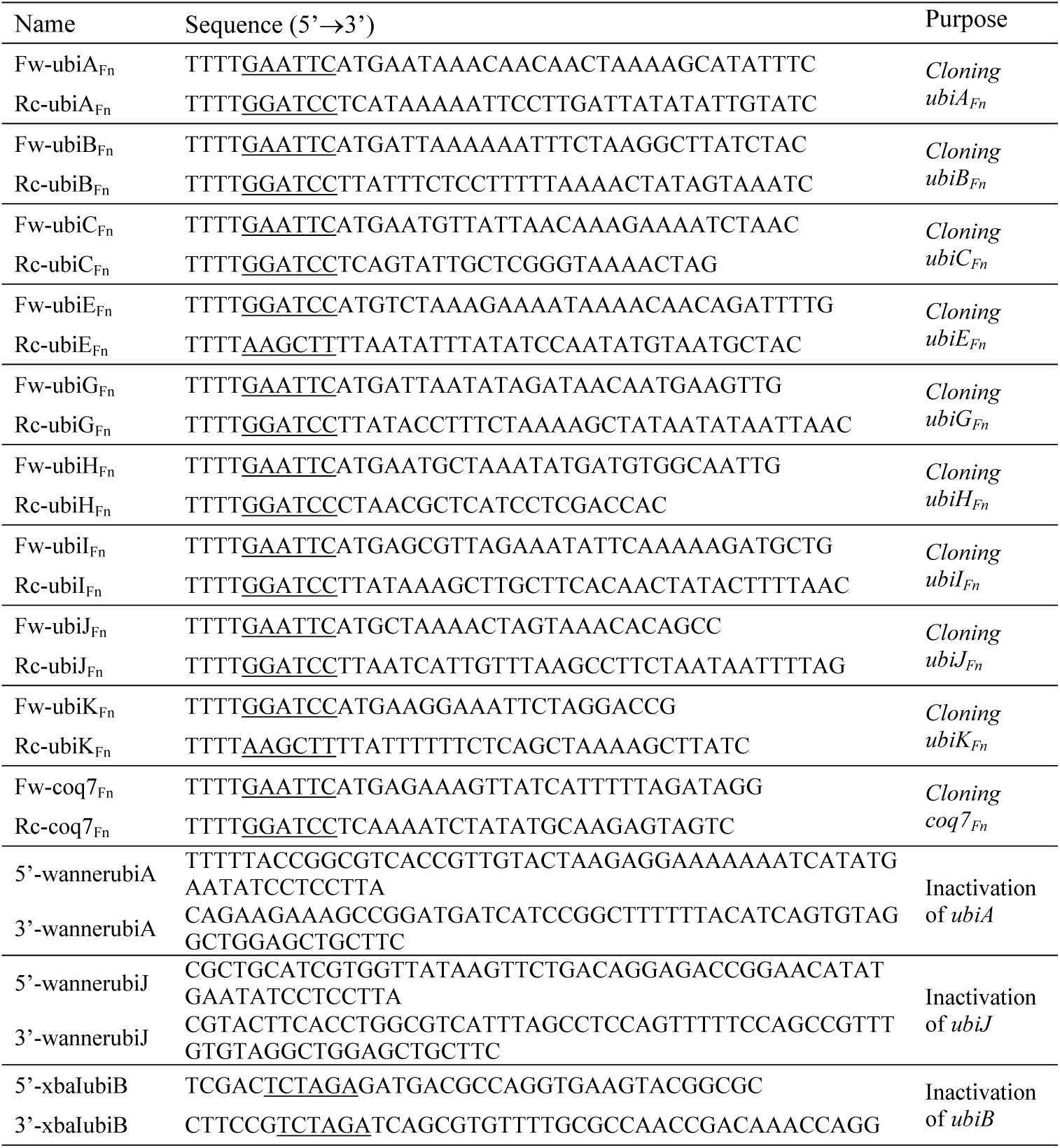
Primers used in this work. Restriction sites are underlined.

